# Two tiers of piRNA clusters balance diversification of piRNAs with limitation of off-target effects

**DOI:** 10.64898/2026.01.28.702291

**Authors:** Axel Poulet, Mariana Witmer, Danyan Li, Romane Cathelin, Josien C. van Wolfswinkel

## Abstract

PIWI-interacting RNAs (piRNAs) are important factors in the protection of the genome against nucleic acid invaders such as transposons. piRNAs are encoded in the genome in regions known as piRNA clusters, but major questions remain surrounding their regulation. Two such questions are how the piRNA clusters are recognized as piRNA sources, and how the piRNA response can expand from these regions to facilitate the recognition of novel invading sequences without risking the targeting of essential cellular mRNAs.

Here, we investigated the piRNA clusters of the planarian *S, mediterranea*, and found that the clusters differ regarding the PIWI proteins their piRNAs bind to, as well as their chromatin features. We uncovered a subset of piRNA clusters that had unique chromatin features and atypical transcription. piRNAs from these clusters were depleted of genic content, stabilized by 3’ methylation and were not dependent on the ping-pong mechanism. We therefore named these clusters Seed clusters. We identified a second set of piRNA clusters that we called Spread clusters, which had features reminiscent of pseudogenes or aberrant genic transcripts. piRNA generation from these clusters relied on ping-pong, and piRNAs largely remained unmethylated. Further, we found that many more regions in the genome generated single ping-pong events, suggesting that ping-pong is used as a means to diversify the collection of piRNA-generating transcripts beyond the Seed clusters.

We propose that this two-tiered organization of the piRNA clusters allows the stable targeting of known genomic threats by the piRNAs generated from the Seed clusters, while the flexible generation of additional diverse piRNAs from Spread clusters as well as other aberrant transcripts increases the sequence space probed by the piRNA system. The absence of methylation on these additional piRNAs decreases their lifespan and limits the chances of a run-away response. This two-tiered organization thus solves a core challenge of the piRNA system and may be a widespread feature of piRNA systems across animals.

## Introduction

piRNAs are important factors in the protection of the cell’s genomic integrity (Malone and Hannon 2009; Ozata et al. 2019). By incomplete sequence complementarity they mark transcripts derived from transposable elements for degradation and thereby restrict the expansion of transposons in the genome (Ozata et al. 2019; Gainetdinov et al. 2023). Further, in some cases piRNAs associate directly with the genomic loci matching these transposons to induce heterochromatin, accomplishing transcriptional silencing at these sites (Brower-Toland et al. 2007; Ashe et al. 2012; Le Thomas et al. 2013; Iwasaki et al. 2016; Li et al. 2021). piRNAs are generated from specialized cellular transcripts that contain fragments of transposon sequences that have previously invaded the organism (Aravin et al. 2007; Brennecke et al. 2007; Duc et al. 2019). This allows the piRNAs to direct silencing to sequences related to these known genomic threats. How the system can also adjust to recognize novel invaders has remained less clear.

Two modes of piRNA biogenesis have been described, colloquially known as “ping-pong” and “primary synthesis” or “phasing” (Aravin et al. 2007; Brennecke et al. 2007; Gunawardane et al. 2007; Han et al. 2015; Mohn et al. 2015; Gainetdinov et al. 2018). Primary synthesis involves the repeated cutting of a single RNA strand into PIWI-bound fragments. In *Drosophila* this can produce head-to-tail (“phased”) piRNAs, but in other systems spacing is more irregular. This biogenesis process requires a single PIWI protein and a nuclease - typically Zucchini. Processing may be triggered by the targeting of the RNA strand by another PIWI-piRNA complex, or by intrinsic properties of the strand. Due to biases of Zucchini and the PIWI protein, the resulting piRNAs mostly start with a 5’ Uridine (Stein et al. 2019). After trimming to the correct size, the piRNAs are methylated on their 3’ end to protect the molecule from further exonuclease activity (Horwich et al. 2007; Kirino and Mourelatos 2007; Saito et al. 2007; Kamminga et al. 2010). In the ping-pong mode, which is generally considered a mechanism of piRNA amplification, two PIWI proteins alternate cleavage of the two complementary RNA strands, thereby repeatedly producing the same pair of methylated piRNAs that have a characteristic 10nt overlap of their 5’ ends, known as the ping-pong signature. Typically one of the piRNAs (initiator piRNA) has a Uridine at the 5’ end (5’U), and the other piRNA (responder piRNA) has an Adenine at the 10^th^ position (10A). Base-pairing needs to be extensive, but does not need to be perfect for the cleavage to proceed (Anzelon et al. 2021; van Wolfswinkel 2023), allowing for a limited amount of drift towards sites with abundant complementary RNA strands. The ping-pong cycle primarily takes place in peri-nuclear RNA-protein condensates known as nuage.

piRNAs are typically concentrated in specific genomic regions, known as piRNA clusters. In *Drosophila*, a handful of clusters produces the majority of germline piRNAs (Brennecke et al. 2007; Gebert et al. 2021). The most prominent *Drosophila* clusters are dual-stranded: piRNAs are generated in both orientations. The HP1 variant Rhino which recognizes the H3K9me3 at these loci mediates this bidirectional transcription specifically in the germline where ping-pong can take place (Klattenhoff et al. 2009; Mohn et al. 2014; Zhang et al. 2014). A few *Drosophila* piRNA clusters generate piRNAs in only one direction, and are labeled unistrand clusters. The best studied example is the flamenco locus (Pelisson et al. 1994; Bergman et al. 2006), which is a specialized region that functions as a transposon trap and by ping-pong-independent piRNA generation protects the somatic follicle cells against active retrotransposons that otherwise could form particles and enter adjacent oocytes (Signor et al. 2023).

Murine piRNAs are all derived from unistrand clusters, but two developmentally distinct populations are generated during sperm development: the pre-pachytene piRNAs and pachytene piRNAs (Aravin et al. 2006; Girard et al. 2006). The pre-pachytene piRNAs largely match transposons. These piRNAs derive from thousands of small genomic loci, and around 50% of pre-pachytene piRNAs come from loci that cannot be defined as a cluster. The pachytene piRNAs overwhelmingly (∼95%) derive from a small number of large clusters, but only ∼50% of the resulting piRNAs match transposons. Thus, while all murine piRNA clusters are unistrand, they still function in different processes and have distinct regulation and biogenesis.

In all other model organisms studied to date, including zebrafish, koala, honeybee, axolotl, and cyclid fish, most piRNA clusters are unistrand (Houwing et al. 2007; Li et al. 2013; Wang et al. 2017; Yu et al. 2019; Konstantinidou et al. 2024; Xiang et al. 2025). In general, they are not as well-defined and transposon-rich as the flamenco locus but can be identified by piRNA density (Rosenkranz et al. 2022; Konstantinidou et al. 2024). It remains unknown whether such piRNA clusters are all equivalent, or whether different types exist, such as in *Drosophila* or in mouse. This distinction is important to allow for analysis of piRNA biogenesis in these models, as well as to understand their dynamics, regulation, and implications for organismal protection. A particular challenge is to understand how animals manage their piRNA populations to be protective but not destructive. As animals frequently encounter new genomic threats (such as newly invading transposons), the piRNA system needs to maintain the flexibility to develop responses to previously unknown elements, which requires plasticity in the response. Increased flexibility however also increases the risk of developing piRNAs that can target coding genes, and will negatively affect the cell. A major question thus is how piRNA-mediated silencing can diversify, but still avoids targeting of essential genes.

Planarians employ an active piRNA pathway in their somatic stem cells, and have developed into a powerful model for the study of piRNA-mediated regulation. The piRNA system of the planarian model *S. mediterranea* consists of two stem cell-specific cytoplasmic PIWI proteins, SMEDWI-1 and SMEDWI-3, and the nuclear PIWI protein SMEDWI-2 that remains present in differentiated cells (Reddien et al. 2005; Palakodeti et al. 2008; Kim et al. 2019; Li et al. 2021; Allikka Parambil et al. 2024). The planarian genome has abundant transposon content, consisting of both DNA transposons and retrotransposons, with several families that remain active to date. The planarian stem cells are located as dispersed cells in the parenchymal space and accumulate at wound sites, where they are easily exposed to novel invaders (Baguna 1976; Salo and Baguna 1984). How planarians balance the protection of their stem cell mRNA transcripts with the need to rapidly recognize transcripts from transposons and newly invading elements is unknown.

Here, we set out to identify the piRNA-generating loci in the *Schmidtea* genome, and find that two different types of piRNA-generating clusters can be distinguished. We identify Seed clusters, that are marked at the chromatin level, and produce the majority of piRNAs, which are stabilized by methylation. In addition, we find a second type of clusters, which we named Spread clusters, that produce diverse, sparse, and largely unmethylated piRNAs, and whose processing is likely triggered by an initial targeting event from a Seed cluster piRNA. The identification of these distinct types of clusters with distinct piRNA products help explain how planarian piRNAs can combine diversification with sequence control.

## Results

### Planarians encode 2 main types of piRNA clusters, based on piRNA content

We designed two complimentary strategies to obtain piRNAs in *S. mediterranea* (**Figure 1A**), and applied these to identify piRNA clusters using a newly designed annotation algorithm (**Suppl Fig 1A;** see Materials and Methods for details). First, we used total small RNA libraries and oxidized libraries (which are enriched in methylated small RNAs that characterize piRNAs) and identified likely piRNAs by read length. We mapped all 28-35nt small RNA reads to the *Schmidtea* genome (**Suppl Fig 1B,C**), and used the uniquely mapping reads supplemented with the multimapping reads to identify piRNA clusters (**Suppl Fig 1A,D**). We identified 16,148 clusters with a median size of 1.9 kb and a median piRNA density of 37.6 TPM. In agreement with previous reports (Friedlander et al. 2009; Konstantinidou et al. 2024), planarian piRNA clusters were largely unidirectional (**Suppl Fig 1E**), and piRNA density was fairly similar over all clusters: no subset of dominant piRNA clusters could be identified (**Suppl Fig 1F**). Clusters were distributed across the chromosomes, and no enrichment at subtelomeric or pericentromeric regions was detected (**Suppl Fig 1G**). The clusters explained 67.5% of the methylated piRNAs, covering 5.4% of the genome (**Suppl Fig 1H**). The remaining piRNAs were spread throughout the genome, and adjusting parameters to increase the included piRNAs to 95% would result in annotating 82% of the genome as piRNA-generating clusters while dramatically reducing piRNA density within those clusters (**Suppl Fig 1D**).

**Figure 1.**
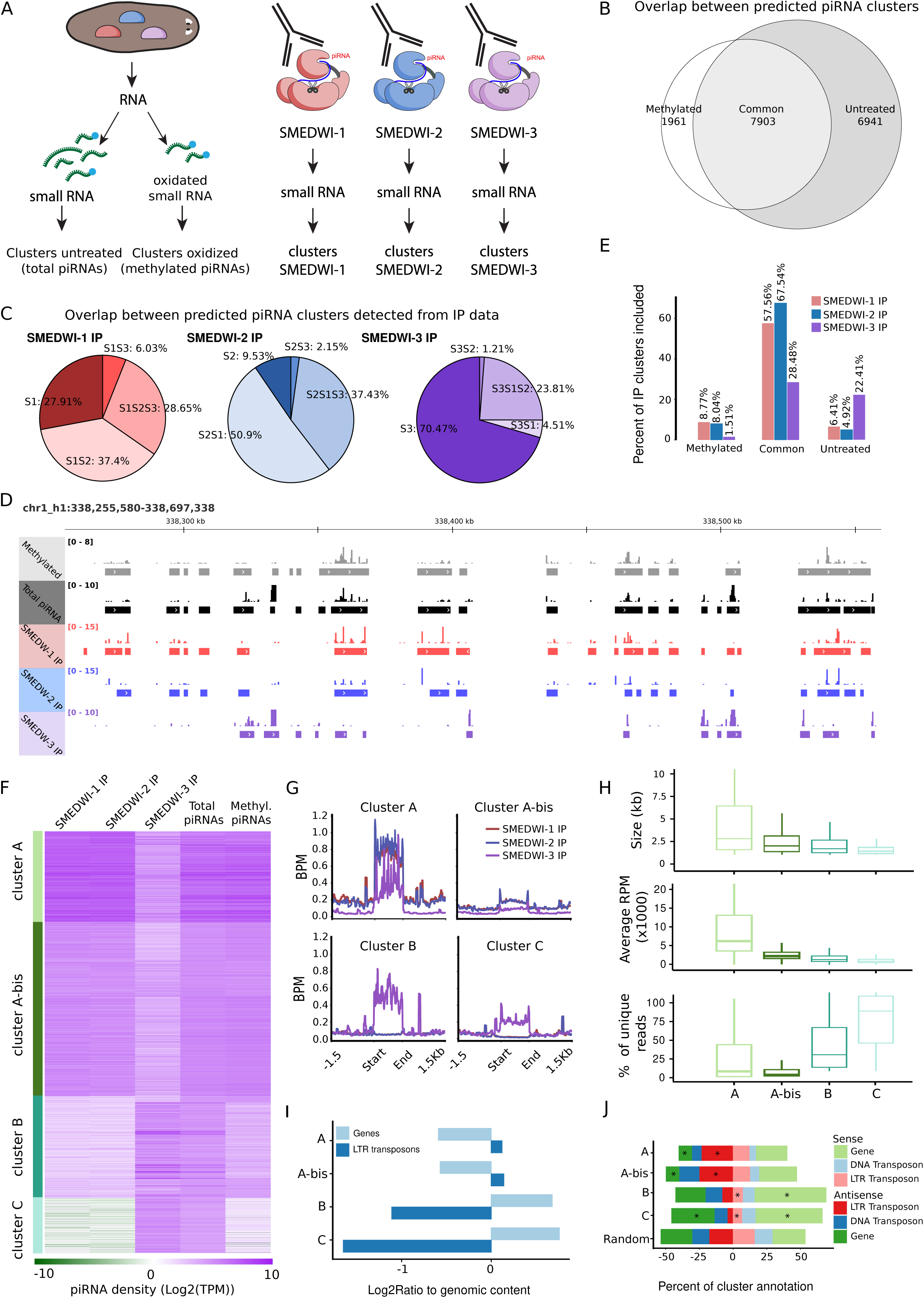
Planarian piRNA clusters separate into 3 major cluster types. **A.** Schematic showing the various strategies used in this study obtain planarian piRNAs and assemble planarian piRNA clusters. **B.** Venn diagram of predicted piRNA clusters based on untreated (total RNAs) or oxidized (methylated RNAs) libraries. The clusters predicted based on total RNA but not based on methylated RNA are referred to as the unmethylated piRNA clusters. **C.** Pie charts indicating the overlaps in piRNA clusters predicted based on piRNAs from individual SMEDWI IPs (Kim et al. 2019). **D.** Genome browser view showing the overlap between the piRNA clusters predicted through the various strategies. piRNA clusters predicted based on methylated piRNAs overlap with clusters based on piRNAs from SMEDWI-1 and SMEDWI-2 IPs, whereas clusters based on piRNAs from SMEDWI-3 IPs are distinct from this set, and rather overlap with clusters predicted solely based on total piRNAs (unmethylated reads). **E.** Bar plot showing the overlap between the clusters predicted based on the SMEDWI IPs and the clusters based on the small RNA isolations. Bars indicate the fraction of the IP-based clusters that falls within the methylated RNA, shared, or unmethylated RNA clusters. **F.** Heatmap of clusters (rows) and piRNA coverage from each of the piRNA libraries, organized by k-means clustering. (k=4; see Supplementary Figure S2A) **G.** Density plots of the piRNA coverage in each of the PIWI IPs for the different cluster types identified in panel F. **H.** Comparison of cluster features (size, read depth, and percentage of unique reads) between the piRNA clusters types. **I.** Enrichment of the cluster content relative to the total genome with regard to genic sequence and sequence related to LTR transposons. **J.** Stacked bar graph showing the fractions of cluster sequence that are annotated as genic or as related to transposon content in each of the two strand orientations. Asterisks indicate notable classes.

Use of untreated (representing total piRNAs) or oxidized (representing methylated piRNAs) small RNA libraries resulted in identification of largely overlapping sets of clusters (**Figure 1B**). Additional clusters found only based on oxidized reads (highly enriched in methylated piRNA) were small and could be detected at sub-threshold read density in the untreated libraries. We therefore considered these clusters to be a subset of the shared clusters. The untreated reads however resulted in the prediction of a significant number of additional clusters that were depleted from the methylated piRNAs (**Figure 1B**), suggesting that these clusters contain largely unmethylated piRNAs, and may represent a substantially different subset of clusters.

As a second, complimentary strategy to identify piRNA clusters, we used small RNAs that co-precipitated with each of the 3 planarian PIWI proteins (Kim et al. 2019) to annotate clusters using the same cluster annotation algorithm. piRNAs bound to SMEDWI-1 and SMEDWI-2 mapped to similar genomic regions, and predicted overlapping piRNA clusters (**Figure 1C, D**). These piRNA clusters were robust as piRNA libraries from different labs predicted the same clusters (**Suppl Figure S1I)**. piRNAs bound to SMEDWI-3 however covered vastly extended genomic terrain and predicted a large set of piRNA clusters that were not identified through SMEDWI-1 or SMEDWI-2. When combining the piRNA clusters based on piRNA-sized small RNA with those based on the PIWI-bound small RNA, we found that the clusters overlapped remarkably well (**Figure 1D, E**). The clusters predicted based on the SMEDWI-1 and SMEDWI-2 bound piRNAs corresponded largely to the clusters that were shared by the oxidized and untreated libraries, whereas clusters predicted based solely on untreated small RNA libraries overlapped mainly with the clusters predicted solely based on the SMEDWI-3 IP.

To uncover patterns within these predicted piRNA clusters, we applied k-means clustering (**Suppl Fig 2A**). We found that planarian piRNA clusters could be divided into 4 types of loci based on their piRNA composition (**Figure 1F,G**). The first two subsets of clusters (type A and type A-bis loci) were dominated by piRNAs bound to SMEDWI-1 and SMEDWI-2. piRNAs from these clusters were particularly prominent in the oxidized small RNA libraries, suggesting that they are (largely) methylated. These clusters explained the majority of all piRNA reads (see also **Figure 3C**). The type A-bis clusters had lower piRNA levels, but otherwise resembled the other type A clusters, and these were therefore grouped together. The second subset of clusters (type C loci) was not covered by reads from the oxidized libraries or from the SMEDWI-1 and SMEDWI-2 IPs, indicating that they relied solely on unmethylated reads that were retrieved in the SMEDWI-3 IP. The third type of piRNA cluster (type B loci) had some coverage from the oxidized libraries and SMEDWI-1 and SMEDWI-2-bound small RNAs, but was still dominated by SMEDWI-3 bound piRNAs (**Fig 1G**). These were therefore considered related to the C-type clusters.

### piRNA cluster types differ in sequence composition

This grouping of the clusters into types held relevance for several other features, including the length of the clusters, the read density, and the mapping frequency of their piRNAs (**Figure 1H**). Type A clusters combined large cluster sizes with high read densities and low fractions of uniquely mapping reads. They contributed the majority of the piRNAs (see also **Figure 3C**) Type C clusters inversely were characterized as small clusters with low read densities and high fractions of uniquely mapping reads.

To investigate the genomic content of the clusters, we annotated the genome with all regions that correspond to transposon sequences as inventoried in Repbase (Bao et al. 2015), and all genic regions as indicated by assembled planarian transcriptomes (see methods). This revealed that type A clusters were enriched in sequence matching transposable elements (mostly LTR transposons) and depleted of genic content relative to the overall genome, whereas type B and C clusters inversely were depleted of transposons and enriched for genic sequences (**Figure 1I, Suppl Figure 2B**). As piRNAs were predominantly generated from one of the two strands of the clusters, we also investigated the orientation of the genomic annotations in relation to the directionality of the piRNAs (**Figure 1J**). Type A clusters were strongly enriched in sequence antisense to LTR transposons, and were strongly depleted for antisense genic annotations. The sense sequence of the A-type clusters was largely unannotated, suggesting that it doesn’t match other genomic elements. Contrarily, type C clusters were strongly depleted of antisense LTR annotations, and were enriched in genic annotations in both directions. Type B clusters were mostly enriched in sense genic annotations, but also contained elevated levels of (mostly sense annotations of) several active DNA transposons including HAT elements, Mariners, and Polintons compared to the rest of the genome (**Suppl Figure 2C**).

Gene-like annotations in the type A clusters were almost exclusively short genes with no introns and low overall expression, suggesting that they may represent accidental bits of Open Reading Frames (ORF) rather than actual genes (**Suppl Figure 2D-E**). The genic annotations in the sense orientation of type B and C clusters had a more typical gene structure, and piRNAs covering the introns of these genic regions were less abundant than those covering the exons and UTRs, suggesting that splicing was completed before piRNA synthesis (**Suppl Figure 2F**). These genic regions however still had low expression levels and small ORFs relative to the transcript length compared to known coding genes, suggesting that they likely represented non-coding transcripts or pseudogenes (**Suppl Fig 2G**).

### Type A piRNA clusters (Seed clusters) have specialized transcription and chromatin markings

The fact that large numbers of piRNAs were derived from specific regions (clusters) indicates that these regions must be transcribed. Indeed, transcripts from the type A clusters could be detected in RNAseq data, and closely matched the regions and direction predicted by the cluster annotation (**Figure 2A,B, Suppl Fig S3**), indicating that the entire transcripts were processed into piRNAs. Transcripts of type B and type C loci could also be detected, but matched their predicted clusters less precisely: transcription tended to occur from wide regions including, but not limited to, the predicted cluster regions. Frequently, the RNA transcription rather matched a transcript annotation that only partially overlapped the clusters, suggesting that only subsections of the B and C type transcripts are processed into piRNAs.

**Figure 2.**
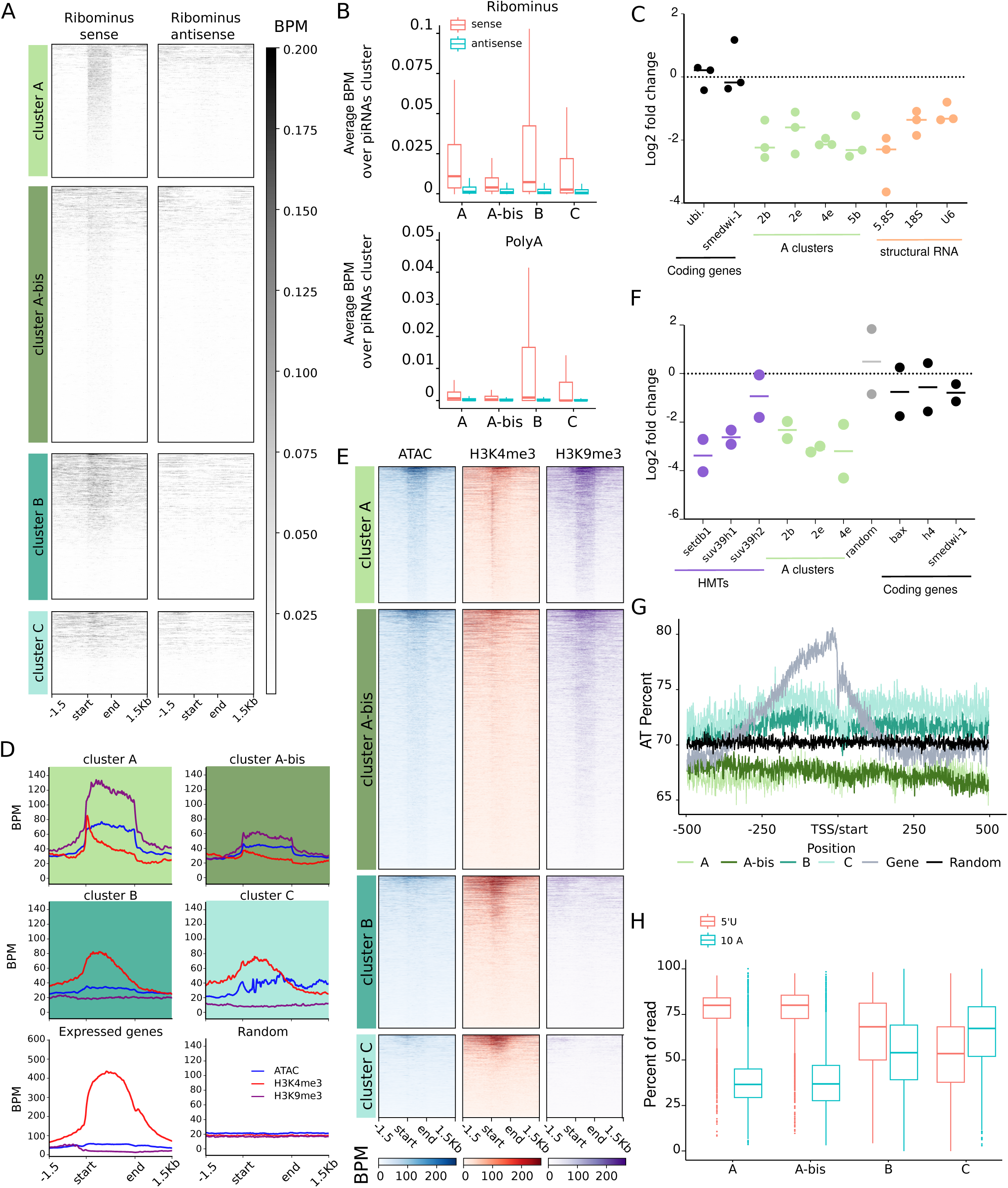
piRNA cluster types differ in their chromatin features and transcription. **A.** Heatmap showing directional ribominus RNAseq read coverage of cluster precursor transcripts for the types of piRNA clusters. **B.** Quantification of sense and antisense reads in bulk RNAseq data for each cluster represented as average bpm over the cluster length), summarized per cluster type. Shown are data from ribominus (left) or polyA (right) RNAseq experiments. **C.** qPCR on total (hexamers) and polyadenylated RNA (oligo dT), demonstrating low polyadenylation on cluster A type transcripts. **D.** Profile plots of ATACseq, H3K4me3, H3K9me3 signal over the piRNA clusters summarized per cluster type, in addition to profiles on genic regions and random genomic regions. Shown are the piRNA cluster regions and the regions 1.5kb upstream and downstream of the clusters. **E.** Heatmap of ATACseq, H3K4me3, H3K9me3 signal around the piRNA clusters (± 1.5 kb). **F.** qPCR comparing levels of piRNA cluster transcripts upon RNAi against the H3K9 histone methyl transferases to control samples, showing decreased transcription in the absence of H3K9me3. **G.** AT-content profile of the Transcriptional Start Sites (TSSs) (±500nt) of the piRNA clusters, in addition to profiles on genic regions and random genomic regions. **H.** Fraction of piRNAs from each cluster that contains a 5’U or a 10A, aggregated per cluster type.

The sequencing data from libraries prepared by polyA selection versus by ribosomal depletion suggested that the transcripts from type A clusters might have reduced levels of poly-adenylation **(Figure 2B**). We therefore tested the adenylation state of the primary cluster transcripts. We found that levels of cluster B transcripts were comparable between cDNA from poly-adenylated RNA and cDNA from total RNA, which is in line with their similarity to mRNAs. Cluster A transcripts however were underrepresented in cDNA from poly-adenylated RNA, indicating that these transcripts frequently lack the polyA tails that are common in genic transcripts **(Figure 2C**). This suggests that the RNA production from A-type clusters is distinct from that of other clusters and of typical mRNAs.

To evaluate how the transcription of the clusters could be regulated, we analyzed the chromatin accessibility around the clusters, as well as the levels of the histone modification H3K4me3 which marks active promoters, and H3K9me3 which marks silenced regions known as heterochromatin. We previously reported that planarian genic regions typically have increased chromatin accessibility over the gene, in combination with a broad peak of H3K4me3 at their 5’ end that extends into the gene (**Figure 2D**) (Poulet et al. 2023). Inspecting the H3K9me3 data, we further found a drop in the level of H3K9me3 at genic loci (**Figure 2D**). The type B and type C clusters roughly followed this pattern although notably weaker. This thus is in agreement with the annotation of these clusters as largely pseudogenic sequences. The type A clusters however had a highly distinct combination of chromatin features (**Figure 2D,E, Suppl Figure S4**): they were clearly more accessible than the surrounding genome, but at the same time had increased H3K9me3 levels. This suggests that these regions are kept accessible, but at the same time are marked for silencing. In addition, the A clusters had a very narrow peak of H3K4me3 at their 5’ end, suggestive of a Transcriptional Start Site (TSS). This, together with the detection of directional long RNAs from these regions, indicates that the primary orientation of the piRNA reads is already determined at the genomic locus rather than being the result of selective stabilization of a subset of (antisense) piRNAs.

We wondered whether the H3K9me3 was required for the transcription from the type A piRNA clusters, or whether (as would be more common) it functioned to reduce transcription from these regions. *S. mediterranea* encodes three H3K9 methyl transferases (in preparation). RNAi-mediated knockdown of these 3 enzymes did not significantly affect transcript levels of several mRNAs or of an unannotated genomic region (**Figure 2F**). Transcripts from the type A clusters however were clearly reduced, suggesting that the H3K9me3 that is found at the A clusters facilitates the production of transcripts that can then be processed into piRNAs.

The type A clusters also stood out on a sequence level: We previously found that genic loci in the planarian genome are characterized by an alteration of the AT content right around the TSS (Poulet et al. 2023): AT content is gradually increased upstream of the TSS, and sharply dips right at the TSS (**Figure 2G**). Type B and type C clusters showed faint signs of a similar sequence organization around their predicted 5’ end. The type A clusters however had lower AT (higher GC) content than random regions, and the GC content increased further into the cluster. This reinforces the notion that the type B and C clusters resemble genic sequences, but that type A cluster loci, while transcribed, are distinct from typical genic loci.

Together, this data indicates that type A piRNA clusters are recognizable as atypical loci by their chromatin and sequence features, likely instructing their transcription and their processing by the piRNA biogenesis machinery. These clusters generate atypical transcripts that lack poly-adenylation, and are processed to form piRNAs from the entire length of the sequence, mostly marked by a 5’U which is a sign of “primary processing” (**Figure 2H**). We thus propose that these clusters should be considered “Seed clusters” for piRNA production. Type B and C loci on the other hand are much harder to distinguish from genic transcripts, although they are low expressed and have limited coding potential. Often only subsections of their transcripts are efficiently processed, and they produce relatively low numbers of piRNAs, many of which have signs of ping-pong activity (**Figure 2H**, and see below). Rather than being specifically regulated, we hypothesize that these loci represent effectively processed targets of the piRNA mechanism. We therefore name the type B and C loci “Spread clusters”.

### 3’ methylation of piRNAs correlates with their PIWI protein but also with their genomic origin

In *Drosophila*, one of the PIWI proteins (Ago3) primarily binds piRNAs of the orientation opposite to the other PIWIs (Brennecke et al. 2007; Li et al. 2009; Sato et al. 2015). Inspection of the orientations of the planarian piRNAs in contrast showed that each of the PIWI proteins predominantly bound piRNAs from the dominant cluster strand (**Figure 3A, Suppl Fig 6A**).

**Figure 3.**
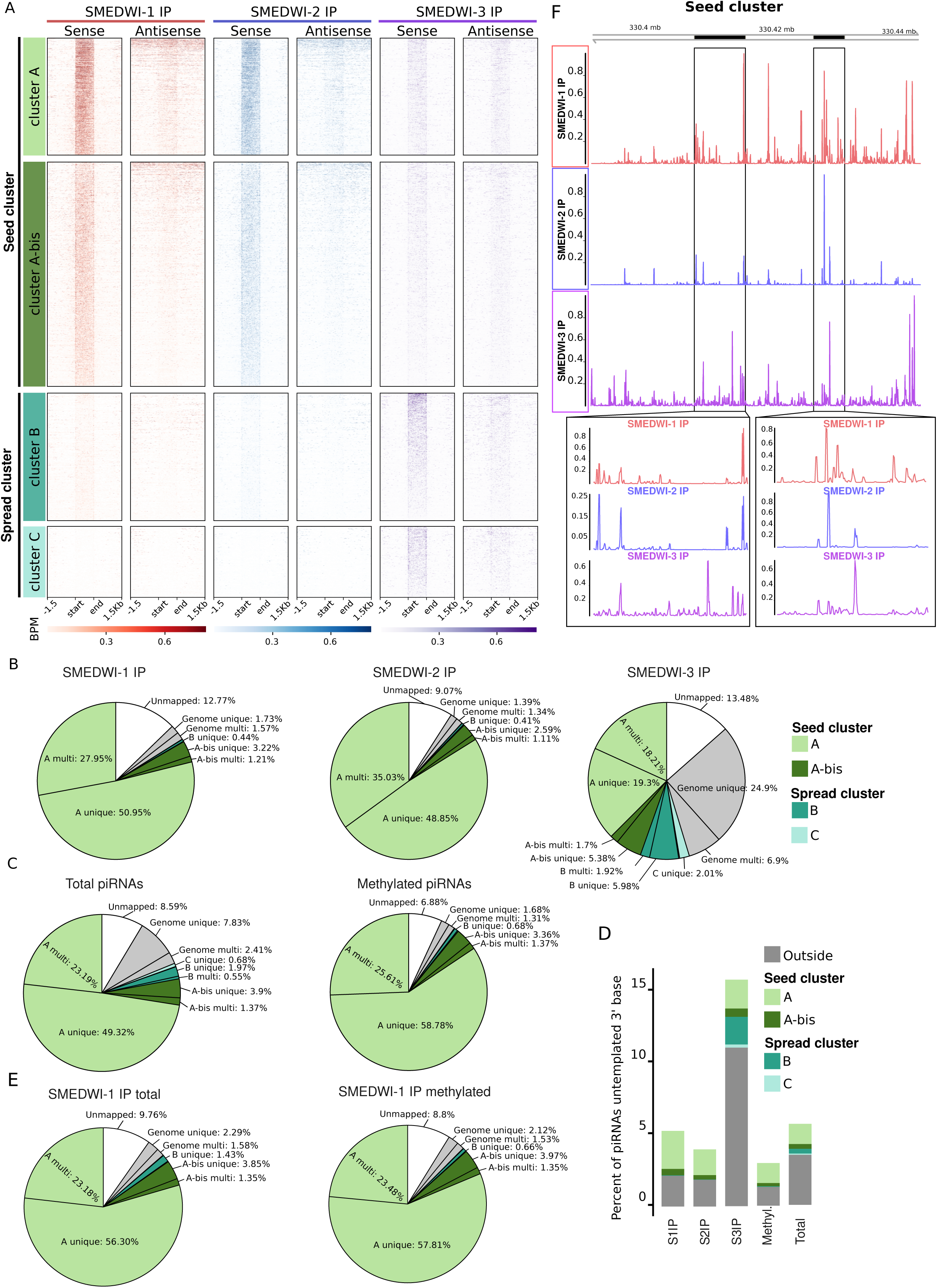
Planarian piRNAs are differentially methylated. **A.** Heatmap showing the coverage and direction of piRNAs bound to each PIWI protein across the different cluster types. **B.** Pie charts showing the contribution of each of the cluster types to the piRNAs bound to each PIWI proteins. **C.** Pie charts showing the contribution of each of the cluster types to the total and methylated piRNA population. **D.** IGV view of an example of cluster A displaying the scale value to the maximum BPM for this cluster for each of the IP of SMEDWI **E.** Percentage of reads with untemplated 3’ extension in each of the small RNA libraries, separated per cluster type. Untemplated bases are more common in SMEDWI-3 bound piRNAs than in piRNAs bound to SMEDWI-1 or SMEDWI-2. Among the SMEDWI-1 and SMEDWI-2 bound piRNAs, untemplated bases are overrepresented in the reads from outside the clusters. **F.** Pie charts showing the mapping of methylated or total piRNAs bound to SMEDWI-1, relative to the clusters types.

The Seed clusters were dominated by piRNAs bound to SMEDWI-1 and SMEDWI-2 (**Figure 1F, 3A**), and inversely, over 80% of the reads bound to SMEDWI-1 and SMEDWI-2 were confined to the predicted piRNA clusters (**Figure 3B,C**). Within the Seed clusters, the reads bound to SMEDWI-1 were consistently more diverse than the reads bound to SMEDWI-2 (**Figure 3F, Suppl Figure S6B,E**): SMEDWI-2 piRNAs were notably more concentrated at discrete positions and these were largely conserved between different experiments. The piRNAs derived from Spread clusters primarily bound to SMEDWI-3 (**Figure 1F, 3A, Suppl Figure S6A-D)**, but an additional 40% of the reads bound by SMEDWI-3 remained unexplained by the clusters (**Figure 3B**). These reads instead mapped sparsely throughout the genome, mostly to non-repetitive sequences. We conclude that SMEDWI-3 broadly binds piRNAs from cluster and non-cluster origin. While the clusters represent genomic transcripts that are effectively processed into piRNAs, SMEDWI-3 additionally binds small RNAs derived from many other loci that are scarcely processed.

The clusters predicted by SMEDWI-3-bound piRNAs overlapped well with the clusters predicted uniquely by unoxidized reads (**Figure 1D,E**), suggesting that SMEDWI-3 piRNAs are likely unmethylated. As an alternative method to estimate piRNA methylation, we investigated the presence of untemplated 3’ bases on the piRNAs (**Figure 3E**) (Li et al. 2005; Ameres et al. 2010; Ji and Chen 2012). Unmethylated piRNAs are more likely to be altered by the addition of untemplated bases, and we indeed found more such events in the untreated libraries than in the oxidized libraries. Further, we found that untemplated bases were overrepresented in libraries generated from SMEDWI-3-bound piRNAs, compared to the other PIWIs, confirming the notion that SMEDWI-3 bound piRNAs have less terminal methylation. Interestingly, when we looked at the genomic origin of the piRNAs with untemplated bases, we found that, independent of the PIWI protein to which they were bound, piRNAs from genomic regions outside of the clusters were overrepresented compared to piRNAs from cluster regions. Further, when we compared untreated libraries from SMEDWI-1 IPs to oxidized libraries, we noted that while both were dominated by piRNAs from the Seed clusters, untreated libraries contained a clear overrepresentation of reads from the type B Spread clusters, and additionally contained higher levels of reads that could not be mapped to the *Schmidtea* genome (**Figure 3F**). This indicates that SMEDWI-1, in addition to the methylated piRNAs from the Seed clusters, also binds piRNAs from the B-type clusters and non-genomic loci, but that these tend to remain unmethylated.

Together, these data indicate that piRNAs bound to SMEDWI-1 and SMEDWI-2 are largely restricted to the Seed clusters and are protected by methylation, whereas the piRNAs bound to SMEDWI-3 derive from a wide range of genomic location and are largely unmethylated. This probably results in a difference in the stability of the piRNAs, as methylated piRNAs are better protected from degradation (Ji and Chen 2012; Gainetdinov et al. 2021). Our data indicate that mainly the piRNAs from the Seed clusters are stabilized by methylation, whereas piRNAs from other locations are probably more easily degraded.

### Ping-pong effectuates diversification of the piRNA response beyond the Seed clusters

To evaluate the contribution of the ping-pong pathway to piRNA generation from the different cluster types, we mapped the occurrence of ping-pong pairs. Ping-pong pairs made up ∼25 % of the total planarian piRNA reads, which is comparable to the fraction in *Drosophila* (∼20 % of total reads). However, when only reads from the oxidized libraries (which contain only the methylated piRNAs) were considered, this fraction was reduced to a mere 6% of reads (**Figure 4A**). This indicates that planarian ping-pong typically involves unmethylated piRNAs on at least one side of the cycle.

**Figure 4.**
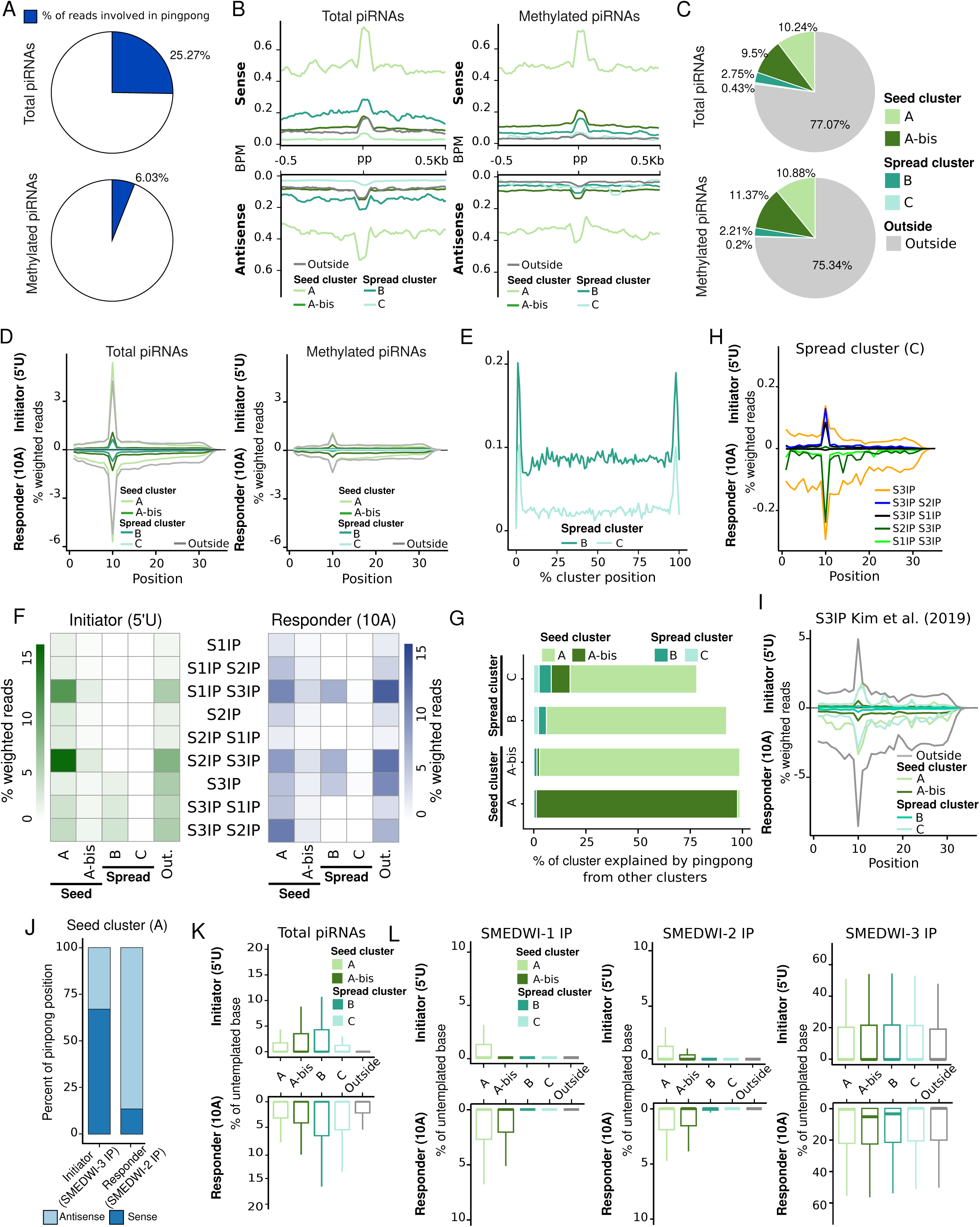
Ping-pong interactions connect the cluster types. **A.** Pie charts showing the fraction of reads among methylated piRNA and all piRNAs for which a ping-pong partner can be identified within the library. **B.** Profile plot showing the piRNA density in regions around ping-pong pairs, aggregated per piRNA cluster type. **C.** Pie chart showing the localization of ping-pong events relative to the clusters, for ping-pong events identified among total piRNAs or among methylated piRNAs. **D.** Percent of weighted reads at ping-pong-related piRNA overlaps, aggregated by the cluster type they derive from, calculated for total piRNAs (left), or methylated piRNAs (right). **E.** Position of ping-pong pairs normalized along the cluster length, showing increased ping-pong at the boundaries of Spread clusters. **F.** Heatmap showing the percentage of weighted reads of the indicated SMEDWI IP involved in ping-pong with the indicated partnering SMEDWI, segregated per cluster type. **G.** Bar graph showing the percentage of each cluster type that contains a targeting site for a piRNA from another cluster type. **H.** Percent of weighted reads at ping-pong-related piRNA overlaps in C-type Spread clusters, separated by PIWI combination of initiator (shown above the axis) and responder (shown below the axis) pair. **I.** Percent of weighted reads at ping-pong-related piRNA overlaps between two SMEDWI-3 bound piRNAs, aggregated by the cluster type they derive from. **J.** Percent of piRNAs that is of sense or antisense orientation relative to the cluster, among the piRNAs involved in ping-pong pairs formed by a 5’U piRNA bound to SMEDWI-3 and a 10A piRNA bound to SMEDWI-2. **K.** Percentage of initiator (5’U) or responder (10A) reads with untemplated 3’ bases in the total piRNA library per cluster, aggregated per cluster type. **L.** Percentage of initiator (5’U) or responder (10A) reads with untemplated 3’ bases in homotypic ping-pong pair, per cluster, aggregated per cluster type.

The ping-pong pathway is generally considered to function in the amplification of piRNAs as loss of ping-pong in *Drosophila* or mouse resulted in a strong reduction of piRNAs (Aravin et al. 2007; Li et al. 2009; De Fazio et al. 2011; Huang et al. 2014). In the planarian clusters however, the read depth of piRNAs that were part of a ping-pong pair was not significantly higher than the read depth of surrounding regions (**Figure 4B**) (beyond what would be expected due to the mandatory presence of a piRNA pair at that site). This indicates that the ping-pong pathway does not accomplish significant amplification or selective stabilization of the two piRNAs involved. The other way by which ping-pong could increase the number of piRNAs is by triggering progressive processing from transcripts that otherwise would not generate piRNAs. We therefore analyzed the genomic locations of the ping-pong events. Surprisingly, when including the complete genome in this analysis, we found that over 75% of the ping-pong positions mapped to genomic regions outside the annotated piRNA clusters (**Figure 4C**). This means that in the majority of the detected ping-pong events, no significant piRNAs were produced from the target RNA except for the direct responder piRNA. It thus appears that transcript cleavage by a PIWI-piRNA complex most of the time does not lead to substantial production of piRNAs from the rest of the transcript, and that the amount of piRNA amplification through the ping-pong pathway is rather limited.

Ping-pong pairs located within the clusters were mostly found in the Seed clusters, resulting in detectable ping-pong signatures (**Figure 4C,D**). In the Spread clusters ping-pong events were more rare and involved lower read counts (**Figure 4D**). This distribution could be expected, as the Seed clusters have higher read density, and are enriched in antisense transposon sequence which can create ping-pong pairs with genomic LTR transcripts. The Spread clusters on the other hand are enriched in sense genic sequence, which makes a ping-pong partner less likely. However, when normalized for piRNA density, ping-pong pairs were actually more frequent in the Spread clusters (**Figure 3C, 4C**), and were concentrated at the outer edges of these clusters (**Figure 4E**). This indicates that piRNA generation from the Spread clusters may well be initiated by ping-pong cleavage, after which most of the piRNAs from these clusters are probably generated by a form of progressive processing that falls outside of the ping-pong pathway.

Together, these findings indicate that ping-pong cleavage is clearly part of the piRNA biogenesis toolkit in planarians, but that the amount of amplification by this pathway is limited. Our analysis of the locations of the ping-pong events indicates that diversification of the piRNA response is a more plausible effect of this pathway, and that the piRNA biogenesis from the Spread clusters may well reflect such diversification.

### Ping-pong responder piRNAs remain largely unmethylated

We used the libraries of SMEDWI-bound piRNAs to determine which PIWI proteins bind to the piRNAs that make up the ping-pong pairs. A previous study had reported that the most common ping-pong combination on transposons was between SMEDWI-2 binding the 5’U initiator piRNA and SMEDWI-3 binding the 10A responder (Kim et al. 2019). When analyzing ping-pong events over the entire genome however we found that the combination of SMEDWI-1 and SMEDWI-3 was equally prominent, but ping-pong pairs involving SMEDWI-1 involved a wider repertoire of piRNAs (**Figure 1C, 3D, 4F**): while SMEDWI-2&3 ping-pong pairs were concentrated on specific regions mostly within the LTR transposon sequences, SMEDWI-1&3 ping-pong pairs were found on more locations within as well as outside of the clusters.

Most ping-pong pairs were located in the Seed clusters, and involved an initiator piRNA bound to SMEDWI-1 or -2, and a responder piRNA bound to SMEDWI-3. Ping-pong pairs located in the type B Spread clusters most commonly involved a SMEDWI-1 or -2 bound piRNA that could have derived from one of the Seed clusters and a SMEDWI-3 bound responder piRNA that uniquely matched the Spread cluster. These ping-pong events were so common that the vast majority of type B Spread clusters could be explained as a target of a piRNA from the Seed clusters (**Figure 4G**). Additionally, homotypic SMEDWI-3 ping-pong occurred frequently in the B-type Spread clusters. Ping-pong pairs in the type C Spread clusters were rare. They occasionally consisted of multimapping reads bound to SMEDWI-1 or -2 that partnered with a uniquely mapping piRNA bound to SMEDWI-3, but more frequently consisted of SMEDWI-3-bound piRNAs on both sides of the cycle (**Figure 4H**). Homotypic SMEDWI-3 ping-pong pairs were less well-defined than those of other PIWI combinations (**Figure 4H,I)**: whereas other pairs showed a strong bias for a 10nt overlap between the partner piRNAs, the SMEDWI-3 pairs formed a wider peak with a long shoulder. This indicates that overlaps of other lengths were also generated, especially when the piRNAs were found in locations outside the clusters (**Figure 4H,I**). This increased flexibility may enable further diversification of the piRNA response. Alternatively, it may indicate that some small regions outside of the clusters are local hotspots for piRNA generation without the involvement of ping-pong.

Ping-pong events initiated by SMEDWI-3 were mostly located in B-type Spread clusters or outside of the clusters (**Figure 4F**). However we found a notable group of ping-pong events between SMEDWI-3 as initiator and SMEDWI-2 as responder on the Seed clusters. In these ping-pong pairs, the majority of the SMEDWI-3 bound piRNAs were of the sense orientation relative to the cluster, and thus produced SMEDWI-2 bound piRNAs antisense to the main cluster transcript (**Figure 4J**). This arrangement thus enables the production of SMEDWI-2-bound piRNAs that could target Seed cluster transcripts, which would be necessary to guide the nuclear SMEDWI-2 to these genomic loci and induce H3K9 methylation.

Most ping-pong responder piRNAs were bound to SMEDWI-3, and thus were likely unmethylated. We indeed found a higher level of untemplated 3’ bases in the responder piRNAs than in the initiator piRNAs in each of the cluster types (**Figure 4K**). To investigate whether the amount of methylation also directly correlated with the position of the piRNA in the ping-pong cycle, we inspected homotypic ping-pong events (**Figure 4L**). In each of the 3 homotypic ping-pong combinations more untemplated bases were found on the responder piRNAs than on the initiator piRNAs, indicating that independent of the PIWI protein, responder piRNAs were less likely to be methylated, and thus are expected to be less stable than the initiator piRNAs.

Together, our data show that the responder piRNAs of the ping-pong mechanism are typically unmethylated. This is established both by their preferential binding to SMEDWI-3, which is highly enriched in unmethylated piRNAs, as well as by reduced methylation of any piRNA in a responder position independent of the PIWI protein to which it is bound. This accomplishes the reduced stability of responder piRNAs, which may be an important mechanism to limit the possibility of off-target effects by these dynamically generated molecules.

### Spread cluster piRNAs, but not Seed cluster piRNAs, depend on SMEDWI-3

To better understand the interactions between the PIWI proteins in the generation of piRNAs, we analyzed piRNAs of PIWI knockdown animals. No major reductions in the overall levels of piRNAs relative to miRNAs were observed in any of the single PIWI knockdowns (**Figure 5A, Suppl Figure S7A**). Loss of SMEDWI-1 resulted in a small decrease in mostly uniquely mapping reads from the Seed clusters, but little effect on the piRNAs from the other clusters (**Figure 5B, Suppl Figure S7B**). As SMEDWI-2 binds similar piRNAs as SMEDWI-1 and remains present, this may be able to largely compensate for the lost SMEDWI-1. Loss of SMEDWI-2 resulted in a similar decrease of total piRNAs from the Seed clusters, but in this case the methylated piRNAs were less affected. Additionally loss of SMEDWI-2 induced a small increase in reads from the Spread clusters (**Figure 5B, Suppl Figure S7B**). This thus confirms the notion that SMEDWI-1 is more active than SMEDWI-2 in the diversification of the piRNAs through unmethylated reads. In agreement with this, we also found that in the *smedwi-2(RNAi)* samples the amount of ping-pong was clearly increased, and primarily involved the Spread clusters (**Figure 5C, Suppl Figure S7F**).

**Figure 5.**
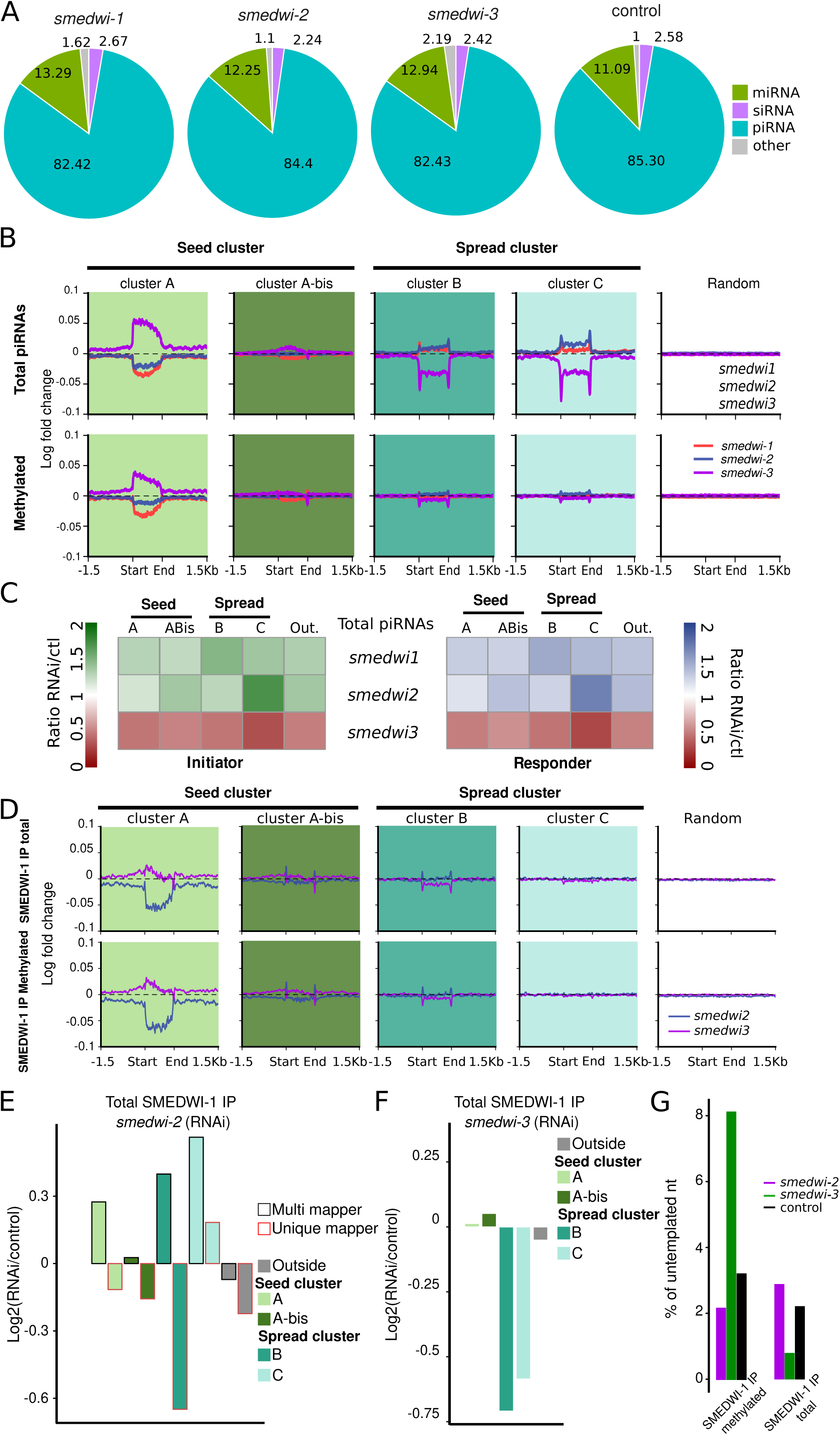
piRNAs from Extender clusters, but not Founder clusters, depend on SMEDWI-3. **A.** Pie charts showing the relative contributions of miRNAs, siRNAs, and piRNAs to the small RNA libraries upon knockdown of each of the Piwi proteins. **B.** Profile plots showing the change in piRNA density (log2 ratio compared to control) around the piRNA clusters upon knockdown of each of the Piwi proteins evaluating total piRNAs or only methylated piRNAs. **C.** Heatmap showing the change in reads involved in ping-pong among total piRNAs upon knockdown of each of the Piwi proteins. **D.** Profile plots showing the change in density of SMEDWI-1-bound piRNAs around the piRNA clusters upon knockdown of *smedwi-2* or *smedwi-3*, evaluating total piRNAs or only methylated piRNAs. **E.** Bar graph showing the log2 fold changes of unique and multi-mapping piRNAs from each of the clusters that are bound to SMEDWI-1 in *smedwi-2(RNAi)* samples versus control samples. **F.** Bar graph showing the log2 fold changes of piRNAs bound to SMEDWI-1 in *smedwi-3(RNAi)* samples versus control samples. **G.** Bar graph showing the percentage of SMEDWI-1-bound piRNAs with untemplated 3’ bases in control samples or upon knockdown of *smedwi-2* or *smedwi-3*.

Loss of SMEDWI-3 expectedly resulted in a notable decrease in total (unmethylated) piRNAs from the Spread clusters (**Figure 5B, Suppl Fig S7A,B**) as well as a strong decrease in the amount of ping-pong (**Figure 5C, Suppl Figure S7F**). Additionally, *smedwi-3(RNAi)* samples had increased piRNAs from the Seed clusters, indicating that the activity of SMEDWI-3 may detract the piRNA biogenesis response from these clusters. These findings are in agreement with the notion that SMEDWI-3 is involved in the diversification of the piRNAs by forming ping-pong pairs into the Spread clusters and the rest of the genome.

To detect the interactions between the PIWI proteins more precisely, we analyzed the effect of SMEDWI-2 and SMEDWI-3 on the piRNAs bound to the neoblast-specific PIWI SMEDWI-1 (**Figure 5D,E**). In the absence of SMEDWI-2, SMEDWI-1 was loaded with less uniquely mapping piRNAs and more piRNAs from repetitive regions (**Figure 5E**), thereby resembling the typical loading of SMEDWI-2. Remarkably, the uniquely mapping piRNAs contribute so much to the normal SMEDWI-1 bound population, that the overall weighted piRNA density from the Seed clusters decreased due to replacement of the unique mappers by the multimapping reads (**Figure 5D**). Multimapping piRNAs were also increased at the Spread clusters at the expense of uniquely mapping reads, but as most SMEDWI-1 bound piRNAs on the Spread clusters are multimappers this did not show in the overall piRNA density at these regions. Nevertheless, these data confirm the notion that the SMEDWI-1-bound piRNA repertoire usually is more diverse than that of SMEDWI-2.

In the absence of SMEDWI-3, no major changes were detected in the Seed cluster piRNAs bound to SMEDWI-1, indicating that the majority of these piRNAs is loaded onto SMEDWI-1 independently of SMEDWI-3 (**Figure 5D,F**). Nevertheless, these piRNAs were more frequently extended by untemplated bases at their 3’ end, indicating that they had remained unprotected for a longer amount of time before methylation took place. This thus suggests that piRNA methylation may be more efficient in the presence of SMEDWI-3 (**Figure 5G**). Spread cluster piRNAs however were lost from SMEDWI-1 upon absence of SMEDWI-3. which was strongest in the total RNA libraries, indicating that these piRNAs were likely unmethylated (**Fig 5D, F, Figure S7E**). This indicates that without SMEDWI-3 less diversification of the SMEDWI-1 piRNA repertoire takes place.

Together, these data indicate that processing of transcripts from the Seed clusters into methylated piRNAs bound to SMEDWI-1 and SMEDWI-2, occurs independently of SMEDWI-3. These Seed piRNAs can target multiple loci, including transcripts from the Spread clusters, which do depend on SMEDWI-3 and remain unmethylated, resulting in a diversification of the piRNA response.

## Discussion

piRNAs cover a large, dynamic, non-random sequence space, and our understanding of the rules for biogenesis of these important regulatory molecules is still incomplete. Several algorithms have been developed to identify regions of high piRNA density on a genome, and this has been used to identify piRNA clusters in several model organisms (Aravin et al. 2007; Friedlander et al. 2009; Lau et al. 2009; Lim et al. 2014; Jehn et al. 2018; Rosenkranz et al. 2022; Konstantinidou et al. 2024). Such clusters likely all represent (parts of) transcripts that are efficiently processed by piRNA machinery, but they may depend on distinct mechanisms for their transcription as well as for their processing into piRNAs. The biogenesis of piRNAs is often discussed in terms of their generation through “ping-pong” or “primary synthesis” / “phasing”. Our findings indicate that while these terms describe the cleavage process that produces the 5’ end of the piRNA, an orthogonal distinction can be made based on the type of cluster transcript that the piRNA is cleaved from. We find that there is no strict correlation between the cleavage mechanism and the type of cluster, and we propose that in terms of their biological impact it may be more important to consider what type of piRNA cluster a piRNA is generated from rather than how it was cleaved.

We combined the identification of piRNA-enriched regions with the analysis of chromatin marks and transcript features, as well as with the analysis of PIWI-bound piRNAs. This revealed that planarian piRNA clusters, which are all unistrand clusters, can be separated into two distinct types of loci. The first type of loci, named Seed clusters, are marked by a special combination of chromatin features: they have H3K9me3 in combination with a sharp peak of H3K4me3 at their 5’ end. They generate primarily non-adenylated transcripts that are processed into methylated 5’ U-biased piRNAs bound primarily by PIWI proteins SMEDWI-1 and SMEDWI-2. Generation of these piRNAs is largely independent of SMEDWI-3. The other type of clusters, named Spread clusters, have chromatin features that resemble those of genic loci. The piRNAs from the Spread clusters are largely unmethylated and bound by the PIWI protein SMEDWI-3. The piRNA production from these clusters may depend on ping-pong targeting by piRNAs from the Seed clusters.

The existence of a subset of clusters that is enriched for the heterochromatic mark H3K9me3, is reminiscent of the dual strand piRNA clusters in *Drosophila.* In *Drosophila*, piRNA generation from these clusters is mediated by the HP1 homologue Rhino, which stimulates bidirectional transcription from these loci. *S. mediterranea* encodes several HP1 homologs, but none with the characteristics of Rhino. Additionally, in contrast to the situation in *Drosophila*, transcription of the planarian Seed piRNA clusters is unidirectional: the piRNAs and long transcripts are strongly biased towards one of the strands, and a peak of H3K4me3 marks the 5’ end of the cluster, suggesting that a single long transcript is likely generated from each cluster. This indicates that planarians must apply a distinct mechanism to transcribe these regions, different from the strategy in *Drosophila*. Our finding that this transcriptional mechanism likely only applies to a subset of the piRNA clusters (namely the Seed clusters) will inform future studies to identify the motifs and molecules involved.

Our data suggest that, at least in planarians, the function of the ping-pong mechanism is the diversification of the piRNA response rather than the amplification of specific piRNA sequences. The piRNAs that make up a ping-pong pair were not significantly more abundant than other piRNAs from the same cluster, arguing against substantial amplification of such pairs by the ping-pong cycle. However, the targeting of a transcript through the ping-pong cycle can induce further generation of piRNAs from this transcript, and thus does result in larger numbers piRNAs - only with sequence different from that of the ping-pong pair. This effect of the ping-pong pathway has been previously proposed (Czech and Hannon 2016; Gainetdinov et al. 2018), but its relative contribution remained unclear. Based on our data, the induction of further processing after a ping-pong hit is relatively rare, as 75% of the ping-pong events occurred outside of clusters, indicating that little further piRNAs were formed from those loci. Nevertheless, we propose that the Spread clusters are the result of efficient piRNA production from some such target transcripts. This thus accomplishes diversification of the piRNA population to include piRNAs with little sequence similarity to the original set of hard encoded piRNA clusters and allows the system to detect a wider range of elements. Why some transcripts are efficiently processed while most ping-pong targeting represents a dead end is not entirely clear from our data. We find that effective piRNA sources have little coding sequence and resemble pseudogenes, and sometimes contain sequence from DNA transposons. The absence of strong ORFs may license these transcripts for piRNA generation (Allikka Parambil et al. 2024). Additionally it is possible that their cellular subcompartment contributes to their selection. Further studies will analyze these aspects in more detail.

The piRNAs bound by SMEDWI-1 and SMEDWI-2, which are largely derived from the Seed clusters, were methylated, and thus protected from degradation. In contrast, the piRNAs bound to SMEDWI-3 did not match the population of methylated piRNAs, and included a high fraction of piRNAs with untemplated 3’ bases, indicating that these piRNAs were not protected by methylation. Interestingly, our protein work on the planarians PIWI proteins frequently only detects fragments of SMEDWI-3 whereas SMEDWI-1 and -2 are routinely detected as full-length proteins, suggesting that the SMEDWI-3 protein is also less stable than the SMEDWI-1 and -2 proteins. This suggests that SMEDWI-3 complexes may have a shorter lifespan than SMEDWI-1 and SMEDWI-2. The implication of this organization may be that the diversified piRNAs bound to SMEDWI-3 are rapidly degraded, limiting their ability to create an avalanche of piRNAs throughout the transcriptome. Further, we noted a specific depletion of sequences matching antisense LTRs from the SMEDWI-3 bound piRNAs (data not shown), which may suggest that SMEDWI-3 piRNAs are subject to a form of Target-Directed Degradation. This would further limit the amount of damage they can do.

We found evidence that besides diversification of the piRNA response, SMEDWI-3 may have an additional role in focusing the methylation of the piRNAs bound to SMEDWI-1 and SMEDWI-2. In the absence of SMEDWI-3, SMEDWI-1-bound piRNAs were more likely to carry untemplated 3’ bases, suggesting that SMEDWI-3 aids in the recruitment of methylation activity to the piRNAs from the Seed clusters. This arrangement may ensure that among the Seed cluster piRNAs, primarily those sequences that have found a target are stabilized and retained for further silencing. piRNAs - from the Seed clusters or from other sources - that have no target and thus only form a risk for deregulation, appear to remain unmethylated for longer timespans, and thus would be degraded more frequently. Further, SMEDWI-3 enabled the generation of SMEDWI-2-bound piRNAs antisense to the Seed clusters, which may help in recruiting H3K9me3 to these loci to retain their distinct chromatin state.

We propose that the uncovered arrangement of two types of piRNA clusters, addresses a central conundrum in piRNA biology, namely the need to efficiently adapt to new threats by diversifying the piRNAs and probing the sequence space, while at the same time limiting the likelihood that cellular mRNAs are swept up in the targeting and become silenced as well. By employing a set of clusters that is enriched for antisense LTR transposon sequence and otherwise largely consists of non-genomic nucleic acid combinations, animals can ensure the basic targeting of known threats without much risk to their own mRNAs and structural RNAs. By then allowing these piRNAs to initiate piRNA generation from other transcripts, the piRNA pool can diversify and explore targeting of even just faintly related transcripts. Limiting the lifespan and/or stability of these downstream piRNAs then reduces the risk of run-away amplification and of targeting mRNAs.

We inspected several other piRNA model systems, but did not find evidence of differential methylation of piRNAs in *Drosophila*, zebrafish, mouse, or hydra. We can only speculate why the planarian piRNA system uses this mechanism to selectively destabilize some piRNAs. It is possible that this is related to the fact that the planarian piRNA system functions in a long-lived cell type, namely the adult pluripotent stem cells. Other piRNA systems, which primarily function in the germline, go through a natural clearing of piRNAs at every generation, as most somatic cells do not have an active piRNA system, and the piRNA population is re-established in the new germline from scratch or from a specifically selected subset of piRNAs. This avoids the uncontrolled diversification of the piRNAs. In planarians, the stem cells, and with it the piRNA system stay present for decades without any such reset. To avoid the diversification to run wild, the rapid degradation of diversified piRNAs might be a necessary control.

Interestingly, while the differential methylation of piRNAs may be specific to planarians, we do find indications that the presence of two distinct types of piRNA clusters can be detected in several other animals. Based on publicly available PIWI IP small RNA data and chromatin data, we find evidence for distinct piRNA cluster types in *Hydra* and among the pre-pachytene clusters in mouse. Combined with the situation in *Drosophila*, which may have a specialized version of the two-type cluster organization, we hypothesize that this system could be widespread and may address a crucial conundrum in piRNA biology, namely the need to efficiently adapt to new threats by diversifying the piRNAs and probing a large sequence space, while at the same time limiting the likelihood that cellular mRNAs are swept up in the targeting and become silenced as well.

## Methods

### Animal husbandry and RNAi treatments

*Schmidtea mediterranea* asexual clonal strain ClW4 was maintained as previously described (Newmark and Sanchez Alvarado 2000). Briefly, animals were cultured in 1× Montjuic salts at 20°C, fed homogenized beef liver paste every 1–2 weeks, and expanded through continuous cycles of amputation or fissioning and regeneration. Animals were starved 1–2 weeks prior to experiments. To facilitate certain RNAi experiments, planarian water was supplemented with Gentamicin (50 μg/ml) to inhibit bacterial growth.

Animals were starved 3 weeks prior to RNAi experiments. RNAi food was prepared by mixing 3 μl of generic food coloring, 2 μl of dsRNA and 50 μl of homogenized beef liver (Rouhana et al. 2013) and fed to animals in 3-day intervals. *smedwi-2(RNAi), smedwi-1(RNAi), smedwi-3(RNAi)*, *unc22(RNAi)* and *suv39h1/suv39h2/setb1 (triple RNAi)* animals were fed over the course of 2 weeks. DsRNA matching *C. elegans* gene *unc-22* was used as a negative control.

### Immunoprecipitation (IP) of SMEDWI-1 and SMEDWI-2

Starting material consisted of 10 million total cells, 1 million neoblasts (X1), 2 million mixed lineage differentiated cells (Xins), 400,000 epidermal cells, 800,000 intestinal cells, or 40 dissected brains from WT animals. Cells were lysed in RIPA lysis buffer supplemented with 5% glycerol, 0.3% Western Blot Buffer (Roche), and protease inhibitor (Roche)). 5% of the lysate was taken as input. Depending on the lysate, 0.25 −1.25 ug of antibody and 5-25ul of Protein A magnetic beads (NEB) were added to the sample. Immunoprecipitation was performed following the NEB protocol. Small RNA was extracted using TRIzol.

### Small RNA library generation

Following TRIzol extraction, IP or input small RNAs were further cleaned up either by eliminating longer RNAs using AmPure (Beckman Coulter), or by selecting for piRNAs with 3′ modifications using NaIO4 mediated oxidation (Vagin et al. 2006). Small RNA libraries were generated following a protocol adapted from the Zamore lab. Briefly, end-blocked, 5′ adenylated 3′ adaptor was ligated using T4 RNA Ligase 2, truncated KQ (NEB). Excess RT primer was added to anneal with the remaining 3′ adaptor, and 5′ adaptor was ligated using T4 RNA Ligase 1 (NEB). Reverse transcription was performed using AMV reverse transcriptase (NEB). PCR amplified libraries 144-158bp in size were selected by PAGE.

### Analysis of small RNA libraries

Reads were filtered and trimmed using fastp (version 0.21.0 with --length_required 18 --average_qual 20). Then sequences were split on function of their size (siRNA and miRNA < 28nt and piRNA ≥ 28nt). We aligned sequences to the reference genomes using STAR (version: 2.7.11a) allowing up to 2 mismatch and 100 alignments per read. Reference genomes used was *S. mediterranea* genome (schMed3, (Ivankovic et al. 2024)). To suppress spliced alignments, we generated genome indexes without annotation GTF files and used the STAR option --alignIntronMax 1.

### Prediction of piRNA clusters by piScan

piScan source code and user manual are available in on GitLab (https://gitlab.com/vanwolfswinkel/piscan). Future versions will be deposited in the same GitLab repository.

*piscan_cluster.py* is a python program for piRNA cluster prediction that takes a coordinate-sorted bam file aligned by STAR as well as the chromosome size file as inputs. First piScan will split the reads mapped in forward or reverse in two different bedgraph, computing the RPM and % of unique reads over a **window size** (default 30nt) defined by the user (Supplemental Figure 1A). The detection will run two times: once for the forward and once for the reverse bedGraph file. To define a cluster, each window is tested for an RPM value ≥ the **RPM (read per million)** (default: unique mapper = 0.05, multi mapper = 0.1) and **% unique mapped reads** ≥ the threshold (default: 1%) a cluster. This cluster is then extended until the length of **empty windows** is > the maximum empty length define by the user (default: 1kb). A window is defined as empty if its RPM is < the RPM threshold used in the previous step. If the potential predicted cluster is ≥ to the minimum cluster size defined by the user (default: 1.2kb)) it is defined as a cluster. Finally, using bedtools intersect (Quinlan 2014), the overlap between clusters detected in forward and reverse orientation is determined. If at least 90% of their lengths overlap, the forward and reverse clusters are merged and defined as bidirectional.

### Analysis of ping-pong signatures

*piscan_pingpong.py* is a python program for piRNA ping-pong prediction that takes a coordinate-sorted bam file aligned by STAR as well as the chromosome size file as inputs. The script defines each position in the genome with at least one read mapped as a potential ping-pong initiator and saves the 5’ coordinate (position 1 of the read). The program then inspects the reads mapped in the opposite direction overlapping this read to determine coordinate of the 5’ end of the responder read. For each initiator coordinate, the responder read coordinate most frequently overlapping with this coordinate is retained.

### RNA-seq library generation

For mRNA-seq libraries of bulk neoblasts and differentiated tissues, RNA was extracted from isolated tissues using TRIzol Reagent (Invitrogen), and libraries were generated using TruSeq RNA Library Prep Kit v2 (Illumina) following manufacturer’s instructions. For generation of RiboMinus libraries, a dedicated planarian probeset (siTOOLs Biotech) was used to deplete RNA samples from rRNA sequences following the manufacturer’s instructions. Remaining RNA was used for library synthesis using the TruSeq RNA Library Prep Kit v2 (Illumina), starting after the polyA selection steps.

### RNAseq analysis

Reads were filtered and trimmed using fastp (version 0.21.0 with --length_required 20 --average_qual 20) (Chen et al. 2018). Datasets were aligned against the *S. mediterranea* genome (schMed3, (Ivankovic et al. 2024)) using STAR (version: 2.7.11a) (--outFilterMismatchNmax 2 - -alignIntronMax 15000 --alignMatesGapMax 15000 --outFilterMultimapNmax 100 --winAnchorMultimapNmax 100) (Dobin et al. 2013). To keep the information of the multi mapper, piscan_cluster.py (default parameters) was used to create bedGraph file used for the profile plot made with deeptools (Ramirez et al. 2014) and compute the average BPM over the piRNAs cluster.

### ATACseq, CUT&Tag and ChIPseq data processing

Reads were filtered and trimmed using fastp ( --length_required 20 --average_qual 20, version 0.23.2) (Chen et al. 2018). Sequencing datasets were aligned against the *S. mediterranea* genome (schMed3, (Ivankovic et al. 2024)) using bowtie2 (-X 2000 –very-sensitive, version 2.5.1 (Langmead and Salzberg 2012). For ATAC and ChIPseq data, duplicate reads were removed using the Picard toolkit (https://broadinstitute.github.io/picard) (MarkDuplicates with REMOVE_DUPLICATES=true). deepTools (version 3.3) was used to obtain bigwig files for ATAC, CUT&Tag (*bamCoverage* --normalizeUsing BPM -bs 5, version 3.5.5). For ChIP seq data bw files were obtained using also deeptools to make the ratio between the IP and the input (*bamCompare* -bs 5 --scaleFactorsMethod SES ). Then *computeMatrix* (scale-regions or reference-point, eeptools, version 3.5.5) was used to obtain the signal of those chromatin data over the piRNAS cluster detected (Ramirez et al. 2014).

### qPCR analysis

Total RNA was isolated by TRIzol (Invitrogen) and quantified by Qubit. To distinguish between changes in global RNA level or in level of polyadenylation, two cDNA preparations were synthesized from the same RNA samples: one primed by hexamers and one primed by oligo dT. For all cDNA preparations ProtoScriptII (NEB) was used according to the manufacturer instructions, using 1 μg RNA as starting material in a 20 μl reaction. cDNA was diluted 1:5 in MilliQ water and 1 μl was applied to a 10 μl qPCR reaction using EvaGreen master mix (Biotium). RT and qPCR reactions of samples and controls were run in parallel in the same plates. qPCRs were run on a QuantStudio 3 instrument (ABI) with the following program: 95°C, 20 s; 40 cycles of 95°C, 5 s; 60°C, 20 s; followed by a melting curve analysis.

## Resource availability

Sequencing data generated over the course of this study has been deposited in the SRA under accession number PRJNA1415216. Public data analyzed in this study were retrieved from PRJNA633618 (Li et al. 2021), PRJNA911251 (Poulet et al. 2023), GSE122199 (Kim et al. 2019), GSE74169 (Duncan et al. 2015), PRJNA982893 (Wiggans et al. 2023).

## Acknowledgements

We thank members of the Van Wolfswinkel lab for support and discussion. We are grateful to the Keck DNA Sequencing Facility at Yale University for services provided. This work was supported by NIH grant R35GM158281 (to JCvW).

## Author contributions

Conceptualization, AP and JCvW.; data acquisition, MW and DL; methodology, AP and JCvW; formal analysis and software, AP and CR; funding acquisition, JCvW; supervision, JCvW.; writing – original draft, JCvW; writing – review and editing, AP and JCvW.

